# Regions of very low H3K27me3 partition the *Drosophila* genome into topological domains

**DOI:** 10.1101/072900

**Authors:** Sherif El-Sharnouby, Bettina Fischer, Jose Paolo Magbanua, Benjamin Umans, Rosalyn Flower, Siew Woh Choo, Steven Russell, Robert White

**Author notes:** Contributed equally (Sherif El-Sharnouby, Bettina Fischer, Jose Paolo Magbanua).

## Abstract

**Background:** It is now well established that eukaryote genomes have a common architectural organization into topologically associated domains (TADs) and evidence is accumulating that this organization plays an important role in gene regulation. However, the mechanisms that partition the genome into TADs and the nature of domain boundaries are still poorly understood.

**Results:** We have investigated boundary regions in the Drosophila genome and find that they can be identified as domains of very low H3K27me3. The genome-wide H3K27me3 profile partitions into two states; very low H3K27me3 identifies Depleted (D) domains that contain housekeeping genes and their regulators such as the histone acetyltransferase-containing NSL complex, whereas domains containing mid-to-high levels of H3K27me3 (Enriched or E domains) are associated with regulated genes, irrespective of whether they are active or inactive. The D domains correlate with the boundaries of TADs and are enriched in a subset of architectural proteins, particularly Chromator, BEAF-32, and Z4/Putzig. However, rather than being clustered at the borders of these domains, these proteins bind throughout the H3K27me3-depleted regions and are much more strongly associated with the transcription start sites of housekeeping genes than with the H3K27me3 domain boundaries.

**Conclusions:** We suggest that the D domain chromatin state, characterised by very low H3K27me3 and established by housekeeping gene regulators, acts to separate topological domains thereby setting up the domain architecture of the genome.

## Background

Our understanding of genome architecture has advanced rapidly in recent years with major insights coming from three approaches; the characterisation of chromatin domains on the basis of their constituent histone marks and associated proteins [1–4], the use of proximity-dependent ligation (3C and derivatives) to define the topology of chromatin within the nucleus [5–9] and the genome-wide mapping of the binding sites of architectural proteins such as the Insulator component, CTCF [10–13]. The combination of these approaches provides a view of the domain organization of the genome with the partitioning of chromatin into topologically associated domains (TADs) linked to the epigenetic landscape of domains of chromatin state and organized by the activities of architectural proteins.

Although the organization of the genome into domains is well established, the processes that form domains and, in particular, the nature of domain boundaries remain unclear. Architectural proteins are enriched at chromatin state domain boundaries; for example CTCF is enriched at the boundaries of H3K27me3 domains [10, 12–14]. Architectural proteins are also enriched at TAD boundaries. In vertebrates, the boundaries of megabase-scale TADs show enriched CTCF binding [5]. Subsequent higher resolution studies revealed a refined TAD map with a median TAD size of 185 kb and an association between orientated CTCF binding sites and TAD boundaries, indicating a link between CTCF-dependent chromatin loops and TAD domains [8, 15]. In *Drosophila*, TAD boundaries are associated with a number of architectural proteins, including several Insulator proteins such as CTCF, BEAF-32, CP190 and Insulator-related proteins such as Chromator (also known as Chriz) [9, 11]. TAD boundaries are characterised by clusters of architectural protein occupancy and boundary strength correlates with the concentration of architectural protein binding [16]. In several systems, mutations affecting the binding sites of architectural proteins have been shown to lead to reorganization of the domain structure with consequences for gene regulation [17–19]. However, architectural protein binding is not the only genomic feature associated with TAD boundaries; other enriched features include gene density, chromatin accessibility, transcriptional activity and housekeeping genes [5, 9, 11]. This raises the question of whether the boundaries are simply interfaces between adjacent TADs or whether they are more complex “boundary regions” with inherent activities and whose chromatin structure may act to separate flanking TADs. An interesting property of TAD organization in the genome is that it appears to be highly consistent between different cell types [5, 8]. Recent studies suggest that the localization of constitutively expressed housekeeping genes in the regions between TADs may provide a basis for a constant pattern of TADs if the constitutively transcribed inter-TAD regions act to demarcate TAD borders [20, 21]. This suggestion also raises questions about the roles of “architectural proteins” such as CTCF, which is implicated in a wide variety of functions. Some appear to be architectural functions, such as its role as a component of Insulator complexes that block enhancer-promoter interaction providing a basis for the establishment of independent regulatory domains in the genome. On the other hand, a significant proportion of CTCF sites occur close to promoters [22] and CTCF has been shown to promote enhancer-promoter interactions [23–25]. Similarly, the *Drosophila* insulator component BEAF-32 predominantly binds close to transcription start sites (TSSs) and regulates gene expression [26], and the insulator component CP190 is strongly associated with actively transcribed genes [14]. Overall it is unclear whether the clusters of architectural proteins at TAD boundaries provide a straightforward architectural function, acting as barriers demarcating the borders of adjacent TADs, or whether these proteins may be more associated with transcriptional regulation of the genes within the boundary regions.

In this paper we present an investigation into boundary regions in the *Drosophila* genome. We find that they can be identified as domains of very low H3K27me3 levels. The genome-wide H3K27me3 profile partitions into two states; very low H3K27me3 identifies domains that are highly enriched in housekeeping genes whereas domains containing regulated genes, irrespective of whether they are active or inactive, contain mid-to-high levels of H3K27me3. The domains depleted in H3K27me3 correlate with the boundaries of TADs and are enriched in architectural proteins, particularly Chromator, BEAF-32 and Z4/Putzig. However, rather than being clustered at the borders of these domains, these proteins bind throughout the H3K27me3-depleted regions and are much more strongly associated with the TSSs of housekeeping genes than with H3K27me3 domain boundaries. We suggest that the primary function of these proteins is linked to the transcriptional regulation of housekeeping genes and that this activity sets up chromatin regions that act to separate topological domains, thereby establishing the domain architecture of the genome.

## Results

### H3K27me3 domains in development

As part of an investigation into changes in the epigenetic landscape during *Drosophila* spermatogenesis, we characterised the profile of the repressive histone mark, H3K27me3, in chromatin from purified primary spermatocytes. The profile is strikingly different to profiles of H3K27me3 found with embryo chromatin. In the embryo data the prominent feature is the enrichment of H3K27me3 in domains containing target sites of Polycomb (Pc) silencing complexes whereas in primary spermatocyte chromatin H3K27me3 appears much more widespread (Fig. 1). Although Pc target domains still show the highest level of H3K27me3 in primary spermatocyte chromatin, much of the genome shows a moderate level of H3K27me3 and domains with a moderate level are clearly distinct from domains where H3K27me3 is very low or absent. Examining other H3K27me3 profiles provides support for a widespread H3K27me3 distribution, for example chromatin from the Kc167 cell line shows significant H3K27me3 in regions matching the moderate domains in primary spermatocytes and also has corresponding regions of very low H3K27me3 (Fig. 1).

**Fig. 1.**
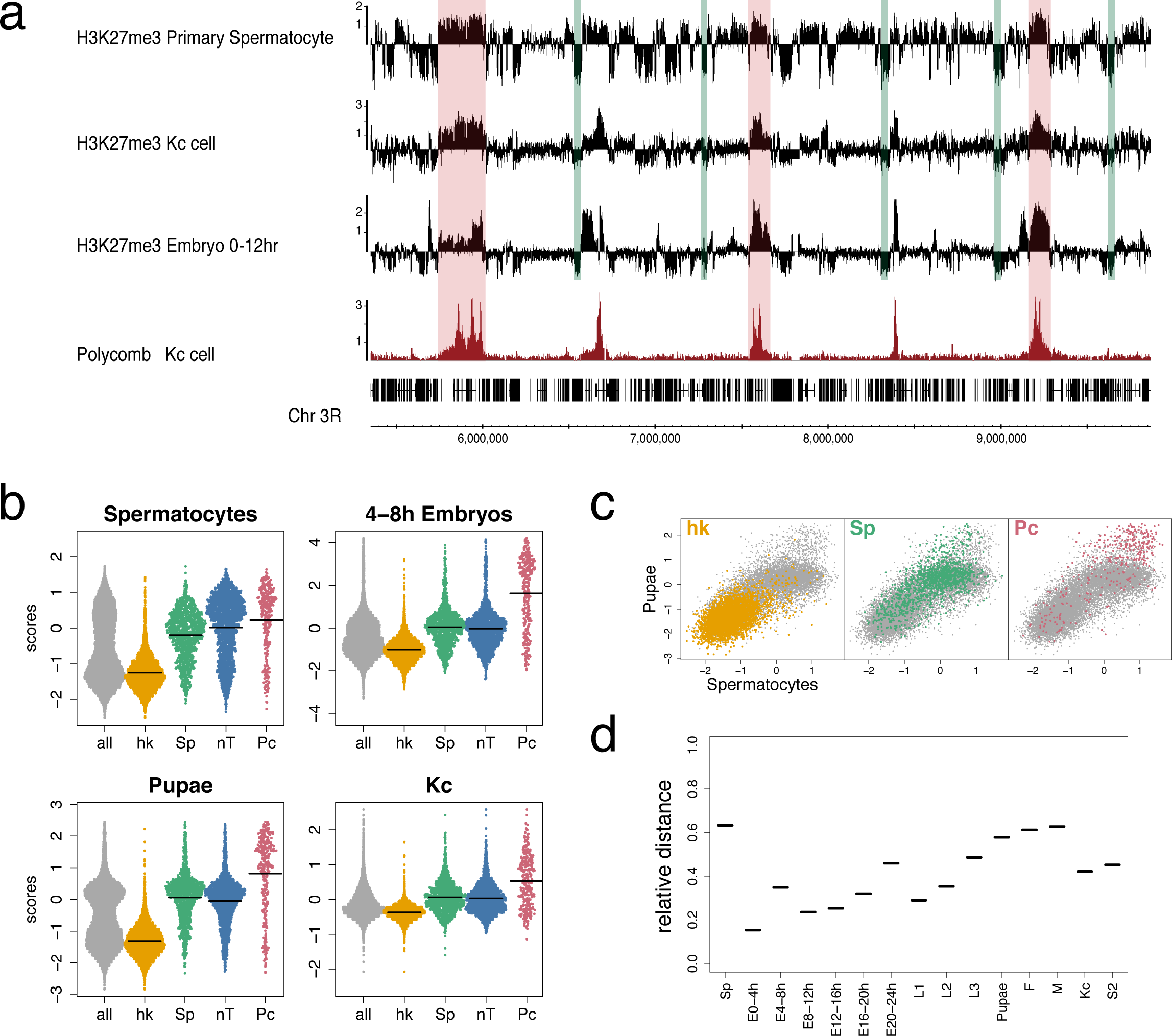
Domains with very low H3K27me3. (a) Binding profiles of H3K27me3 in primary spermatocytes compared to Kc cell and 0-12 hr embryo H3K27me3 profiles and Kc cell Polycomb profile. Green bars indicate selected regions of very low H3K27me3 and red bars indicate selected Polycomb binding regions. See Table S2 for data sources. (b) SinaPlots [69] of H3K27me3 median binding scores at the TSSs (+/-500bp) of different gene classes: all = all genes in euchromatic genome, hk = housekeeping genes, Sp = spermatogenesis genes, nT = genes not expressed in testis, Pc = Polycomb targets from [61], see Methods for derivation of gene classes. Means are indicated by horizontal lines. (c) Scatter plot of H3K27me3 median binding scores at all TSSs (+/- 500bp) for pupae versus primary spermatocytes showing distinct clustering of the different gene classes. (d) Relative distance of the mean of the spermatogenesis gene class binding scores at TSSs, in reference to the hk mean as zero and the Pc mean as 1, at different developmental stages (E = Embryo, L1-3 = Larval instars, F = adult female and M = adult male), in cell lines (Kc and S2) and in primary spermatocytes (Sp). There is a trend of increasing H3K27me3 during development.

To associate levels of H3K27me3 with states of gene expression in primary spermatocytes, we examined the H3K27me3 levels at the TSSs of different gene classes. We find that the lowest levels of H3K27me3 are tightly associated with constitutively expressed (housekeeping) genes (Fig. 1b). In contrast, regulated genes, whether active or inactive, are associated with higher levels of H3K27me3. Induced genes active in primary spermatocytes (spermatogenesis genes) have moderate levels of H3K27me3, whereas inactive genes (not expressed in testis) and Pc targets are associated with higher H3K27me3 levels. In other cells it is also clear that significant levels of H3K27me3 are not only associated with canonical Pc target genes (Fig. 1b,c). A developmental time course indicates that whereas housekeeping genes are always associated with very little or no H3K27me3, the “moderate” class represented by regulated genes (spermatogenesis and not expressed in testis classes) shows increasing H3K27me3 levels. Whilst significantly higher than the housekeeping levels in all profiles, the “moderate” levels are relatively low early in development but then rise to become closer to the level associated with Pc-targets by the pupal stage (Fig. 1d). This suggests that, apart from the housekeeping genes, there is a general developmental increase in H3K27me3 levels outside Pc domains, emphasizing the difference between constitutively active housekeeping regions and the rest of the genome.

### H3K27me3 domains provide a binary partitioning of the genome

Focusing on the domains with very little or no H3K27me3 we investigated their associated genomic features (Fig. 2). H3K27me2 exhibits a similar profile to H3K27me3 with a widespread distribution that has been associated with a global repressive role [27], and we also observe domains of low signal corresponding to the low level domains of H3K27me3. H3K27me3 and H3K27me2 differ at canonical Pc-targets which show high levels of H3K27me3 but low levels of H3K27me2 [27]. The H3K27me3 low level domains also show a link to histone acetylation with a striking match to regions of Histone deacetylase 3 (HDAC3) binding and also a correspondence with regions enriched for MRG15 (a component of the Tip60 Histone acetyl-transferase (HAT) complex, [28, 29]), MBD-R2 (a component of the NSL complex which regulates housekeeping gene expression and is associated with the HAT Males absent on the first (MOF) [30–33] and also the H4K16ac modification, generated by MOF and regulated by HDAC3 [34, 35].

**Fig. 2.**
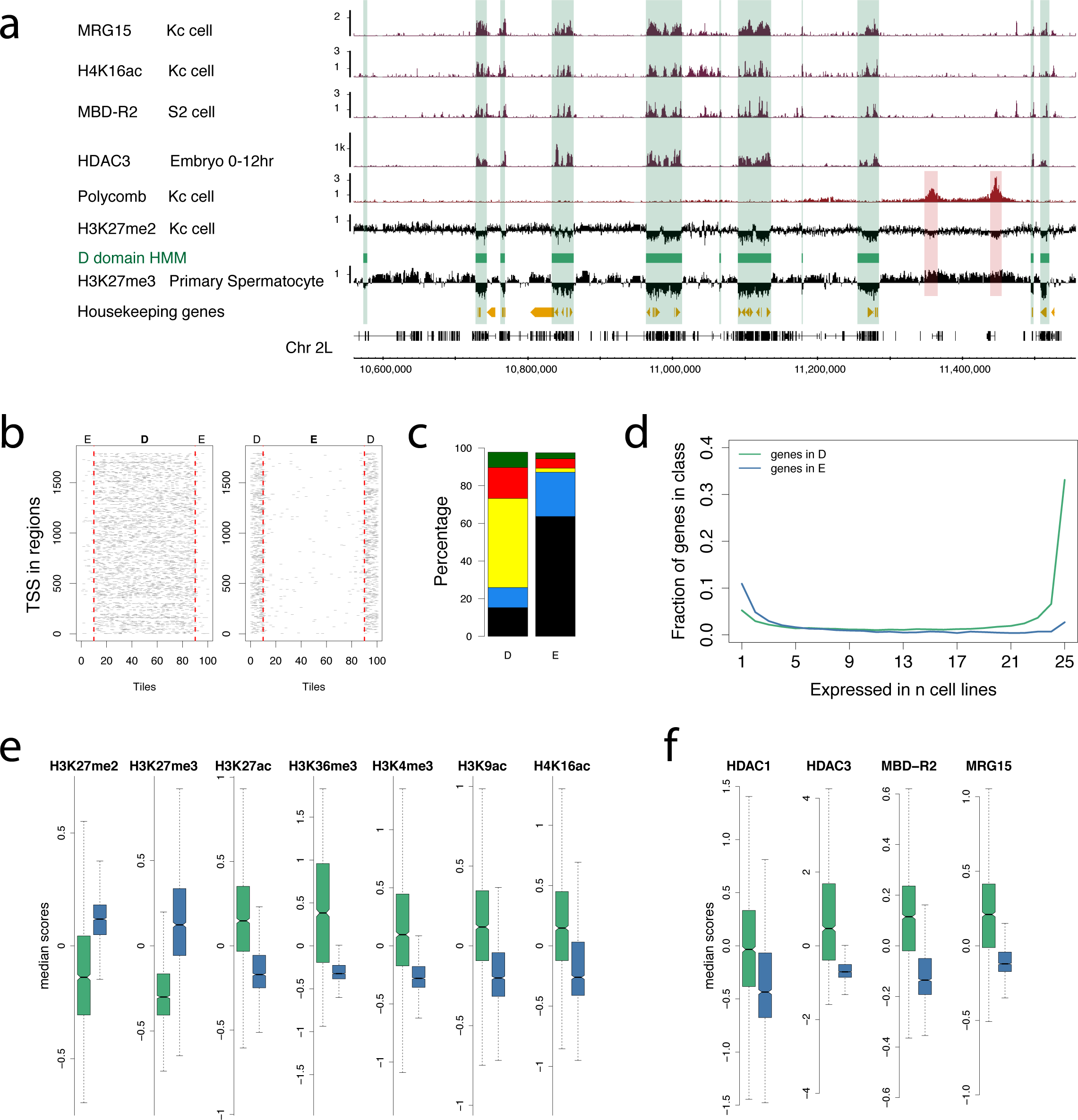
Genomic features of D domains. (a) Profiles of selected histone modifications, chromatin components and housekeeping genes show association with the D domain HMM state (green bars) based on the H3K27me3 profile in primary spermatocytes. The H3K27me3 and H3K27me2 profiles both show low levels in the D domains but differ in the Pc target regions (red bars). (b) Stacked plots of scaled D domains (left) and E domains (right) showing enrichment of tiles containing housekeeping gene TSSs in D domains. D and E regions are split into 80 equal sized tiles per region and extended on both sides by 10 tiles outside the region. Red dotted lines show domain borders. (c) Using the 5-state colour-coded chromatin state classification of Filion et al. [1], the plot shows the different chromatin state proportions in D and E domains and the prevalence of the “yellow” state in D domains. (d) Using cell line gene expression data from Cherbas et al. [68], the plot shows that genes with narrow tissue expression are more prevalent in E domains (blue), whereas genes with broad expression are more prevalent in D domains (green). (e) Boxplots of ChIP scores for histone modifications across D (green) and E (blue) domains showing depletion of repressive chromatin modifications and enrichment of active chromatin modification in D domains. (f) Boxplots of ChIP scores for selected chromatin components showing enrichment in D domains. See Table S2 for data sources.

Applying a Hidden Markov Model to the primary spermatocyte H3K27me3 profile we partitioned the genome into two states, Depleted (D) and Enriched (E) domains (Fig. 2) to facilitate a quantitative analysis of genomic features associated with these regions. There are 1,795 D regions and 1,796 E regions encompassing 34% and 65% of the euchromatic genome respectively. The D regions contain 62% of TSSs and are strongly enriched in housekeeping genes: containing 95% of TSSs of genes in our housekeeping gene class. In contrast, regulated gene classes are associated with E domains. For example, genes whose expression is limited to only a few *Drosophila* cell lines are predominantly in E domains (Fig. 2d). In addition, 75% of embryo Pc targets and 80% of genes specifically activated in spermatocytes are in E domains. Using the five chromatin state classification from Filion et al. [1], we find that D regions are also enriched in active chromatin states, particularly YELLOW which has previously been associated with housekeeping genes and the Non-Specific Lethal (NSL) complex [32]. While the D regions are depleted in the H3K27me3 and H3K27me2 repressive histone marks, they are enriched for a variety of active marks including H3K4me3, H3K9ac, H3K27ac, H3K36me3 and H4K16ac. Binding of the HAT associated proteins MBD-R2 and MRG15 is enriched in D regions along with HDAC1 and HDAC3. Overall, the distinctive features of D and E domains support the potential functional relevance of this binary partitioning of the genome.

### D and E domains and genome architecture

To explore the relationship between the organization of D and E domains and the overall domain architecture of the genome, we generated a high resolution interaction map of the Kc167 cell genome using HiC [6, 36] with fragmentation using the 4-base cutter DpnII. As in previous studies, the interaction map shows the organization of the genome into a series of TADs allowing the derivation of a set of TAD boundaries across the genome. The TAD boundaries show a striking coincidence with D/E domain boundaries with 78% of TAD boundaries occurring within one 10kb bin of a D/E boundary (Fig. 3). A similar correspondence is also found with the TAD boundaries defined in embryo chromatin [9]. Comparing the interaction map with the organization of the genome into D and E domains reveals a clear connection, with the prominent interaction-dense TADs generally corresponding to E domains and intervening regions corresponding to D domains (Fig. 3a, Fig. S1). This correspondence has implications for the interpretation of the overall TAD architecture as it suggests that prominent TADs do not simply abut each other with a discrete boundary at the junction. Rather, it suggests a model where neighbouring prominent TADs are separated from each other by an intervening region corresponding to a D domain (Fig. 3c).

**Fig. 3.**
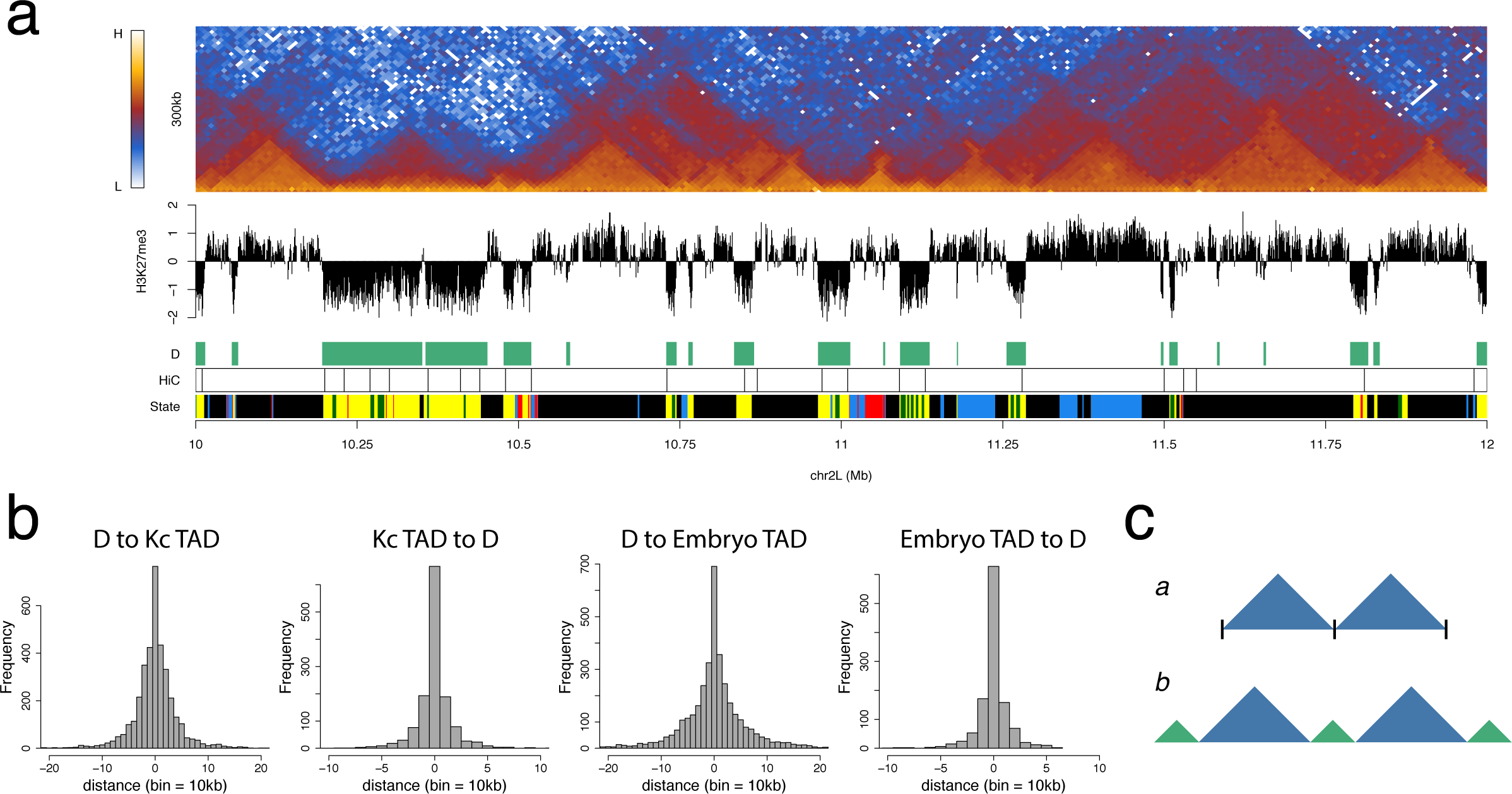
D domains are strongly associated with TAD architecture. (a) Heatmap of 10kb binned normalised HiC interactions across a 2 Mb region of chromosome 2L in Kc cells showing the association of the D domain state and TAD architecture. Below the interaction heatmap the tracks are “H3K27me3” showing the H3K27me3 primary spermatocyte profile, “D” showing the D HMM-defined domains, “HiC” showing the TAD boundaries derived from the Kc cell HiC data and “State” showing the 5-colour chromatin state profile. (b) Histograms showing (from the left) the mapping of D domain borders to Kc HiC TAD boundaries (D to Kc TAD) and vice versa (Kc TAD to D), followed by the mapping of the D domains borders to the TAD boundaries identified in embryo chromatin by Sexton et al. [9] (D to Embryo TAD) and vice versa (Embryo TAD to D). (c) In contrast to model “*a*” where TADs abut at simple interfaces, we suggest model “*b*” where prominent TADs (blue) are separated by boundary regions corresponding to D domains (green).

### D domains are rich in a subset of insulator components

The boundaries between TADs are enriched in insulator protein binding; this is seen for CTCF in vertebrates [5, 8, 15] and for a number of insulator components in *Drosophila* [9, 11, 13, 16]. We analysed insulator component binding in D domains, using published profiles (Fig. 4). We find a strong association between the binding of the insulator protein BEAF-32 and D domains, with 86% of BEAF-32 binding sites mapping to D domains. BEAF-32 binding is known to be largely overlapping with the insulator-related protein Chromator [37], the factor most strongly associated with TAD boundaries in embryo chromatin [9] and which directly interacts with BEAF-32 to form a complex that recruits CP190 [37]. We find Chromator is also strongly enriched in D domains, with 88% of sites mapping to D regions. Chromator forms a complex with Z4/Putzig that recruits the JIL-1 kinase to promote H3S10 phosphorylation [38, 39] and we find Z4/Putzig binding almost entirely confined to D domains with 96% of sites mapping to D regions. Whilst BEAF-32, Chromator and Z4/Putzig form a strongly D-associated group, the insulator component CP190, which is recruited to chromatin by several distinct DNA-binding insulator proteins including BEAF-32, CTCF and Su(Hw), is more widely distributed with only 66% of binding sites in D regions. As over 70% of CP190 sites within D regions overlap with BEAF-32 binding, this suggests that the CP190 in D domains is specifically recruited into BEAF-32 insulator complexes. The insulator components GAF and CTCF are also widely distributed and lack a strong association with D domains. The Su(Hw) insulator protein has a distinct distribution, being relatively depleted from D domains, with only 26% of sites in D regions.

**Fig. 4.**
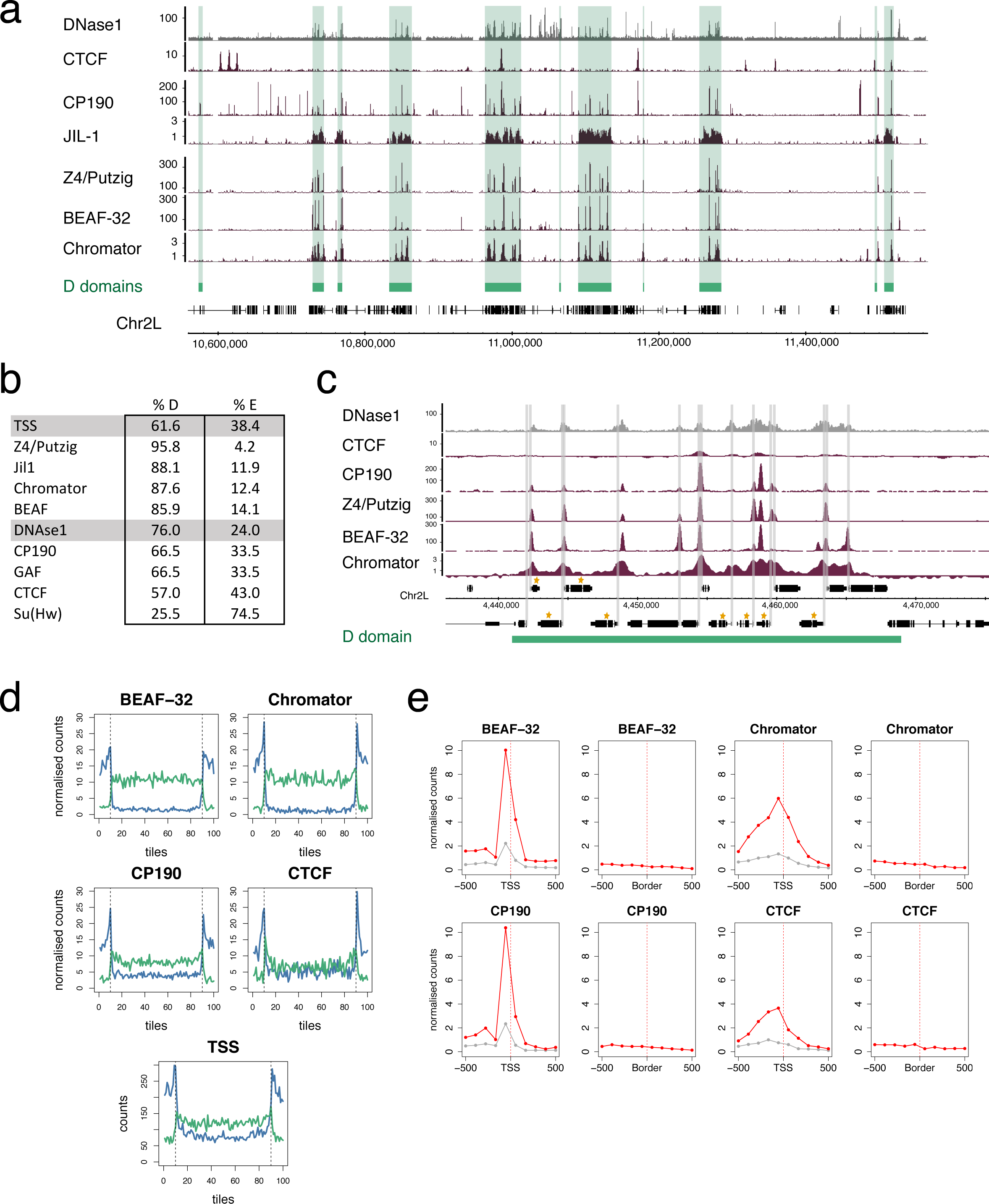
D domains and Insulators. (a) Binding profiles of insulator and insulator-related components showing that a subset, Chromator, BEAF-32, Z4/Putzig and JIL-1, show a clear association with D domains (green bars) whereas CTCF does not. All profiles are from Kc cells, see Table S2 for data sources. (b) Percentages of TSSs, DNase1 sites, insulator and insulator-related components mapping to D or E domains. (c) Profiles of insulator/insulator-related components within a single D domain. The binding sites of these components are spread throughout the domain and show an association with TSSs and not with the domain boundaries. The extent of the D domain is represented by the green bar, TSSs are indicated by grey bars and genes in the housekeeping gene list are indicated by gold asterisks. (d) Plots showing mapping of insulator/insulator-related components to scaled D (green line) and E (blue line) domains. D and E domains are split into 80 tiles per domain and extended on both sides by 10 tiles. The number of normalized binding summits were counted for each tile. For normalisation, the summit counts were scaled by the number of summit binding positions (n) for each factor so that each summit count is 1000/n. The dotted vertical lines indicate the domain borders. For the TSS plot, “counts” is the number of TSSs per tile. The binding locations are evenly spread throughout D domains and not concentrated at the domain borders. For E domains, the flanking peaks are likely due to the scaling causing compression of large domains and a similar profile is seen with insulator/insulator-related and TSS mapping. (e) Accumulation plots focused on either TSSs or domain borders. In the TSS plots the red line indicates housekeeping genes and the grey line indicates non-housekeeping genes. The total summit counts for each factor are scaled as in (d), normalised to correct for the different numbers of TSSs and Borders and the final values scaled to give a range between 0-10.

Although insulators have previously been associated with the borders of chromatin state domains, it is striking that the D domain-associated components BEAF-32, Chromator and Z4/Putzig are distributed quite evenly across D regions rather than being concentrated at the D/E boundaries (Fig. 4c and d). This argues against the idea that insulator complexes are simply positioned at the boundaries of chromatin state domains.

As several *Drosophila* insulator proteins have previously been shown to be enriched at promoters, we examined the association between architectural protein binding and housekeeping gene TSSs. BEAF-32, Chromator, CP190 and Z4/Putzig all show strong enrichment at housekeeping TSSs. Comparing the enrichments at housekeeping TSSs versus D/E borders we find much more enrichment at TSSs than borders. CTCF shows less strong enrichment at housekeeping TSSs, but interestingly it is still more strongly enriched at housekeeping TSSs than at D/E domain borders (Fig. 4e).

### D and E domains are topologically distinct

As noted above, the E regions generally correspond to prominent TADs, i.e. regions of enhanced interaction with widespread interactions across the domains (Fig. 3). In contrast, D domains, even if they are long, have a different appearance in the interaction map with less prominent long-range interactions (Fig. 5). This difference is evident in a plot of short-range (5-50kb) versus long-range (50–500kb) interactions where the D domains correspond to regions with high short-range versus long-range interaction ratios. Accumulating the interaction length data across the genome, we find that D and E domains have significantly different profiles (Fig. 5b) with D domains associated predominantly with short-range interactions and a more rapid fall in interaction density with increasing interaction length.

**Fig. 5.**
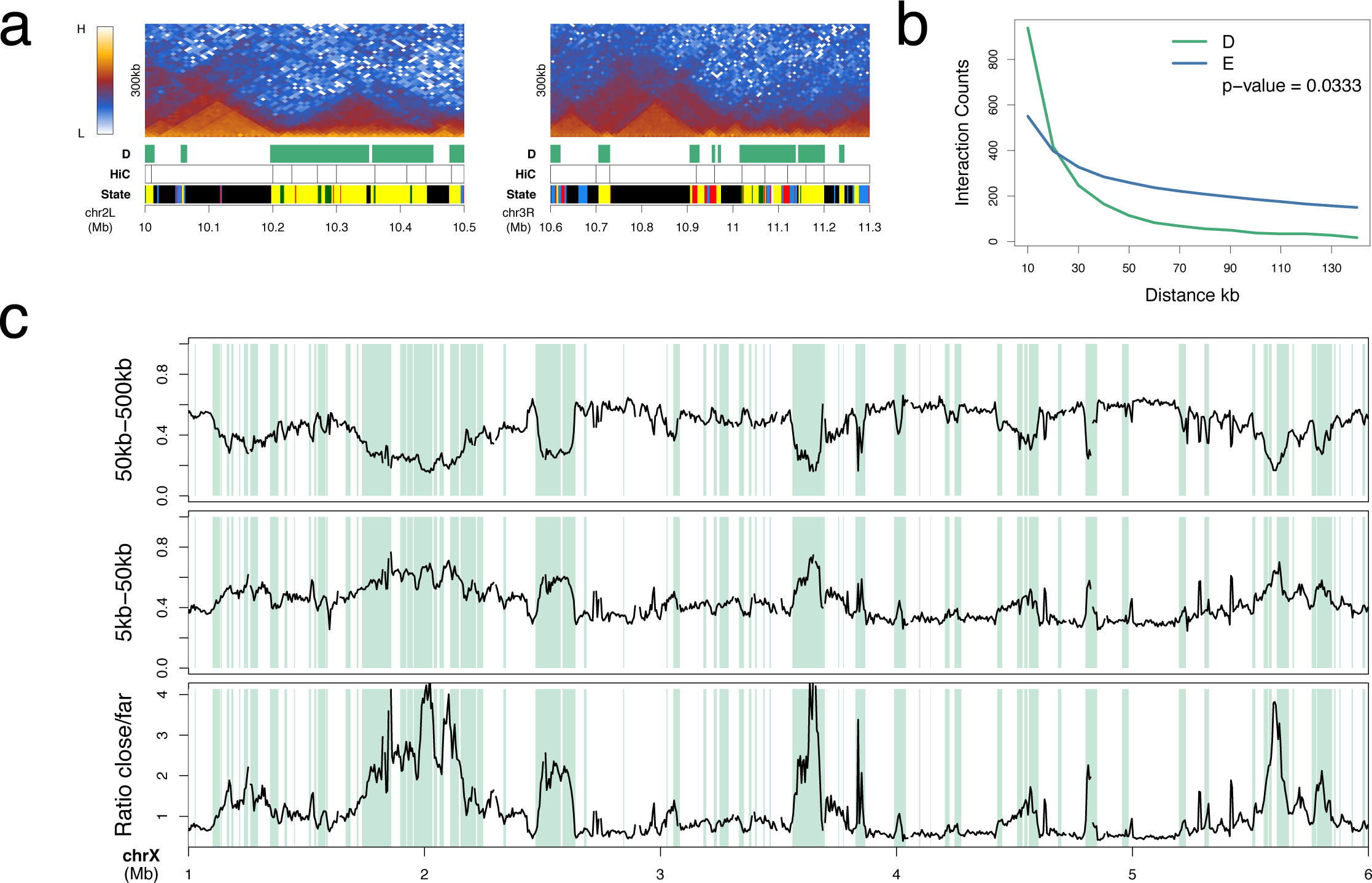
D and E domains are topologically distinct. (a) Kc Hi-C interaction heatmaps at selected D and E domains with tracks “D” showing the D domain as green bars, “HiC” showing the derived TAD boundaries and “State” showing the 5-colour chromatin state profile. Whilst the E region TADs show rather uniform interaction across the whole domain, in the D regions there is a prevalence of short interactions with high heatmap intensity close to the baseline. (b) Interaction distance profile for D domains (green line) and E domains (blue line) showing the preponderance of short interactions in D domains. For both D and E regions longer than 30kb (D = 443 regions, E = 755 regions) the normalised HiC interaction counts (10kb resolution) were collected into 10kb distance bins and the median per bin calculated. The p-value was calculated using the Wilcox test of the medians between D and E. (c) Profiles of interaction distance across a 5 Mb region on Chromosome X (see Methods for details). D domains are indicated by green bars. The “far” interaction (50-500kb) frequency dips in D domains (top profile), whilst the “close” interaction (5-50kb) frequency peaks in D domains. The bottom profile shows the ratio of “close”/“far” interactions.

### Interaction of D domains

In the genomic interaction maps there is evidence for interaction between D regions. This is seen as areas of enhanced interaction that sit on top of E TADs, corresponding to interaction between the two D domains that flank the E TAD (Figs. 1 and 6). Although they are not present at all E domains, such D-D interaction regions are frequently observed. For example, if we take the 174 E domains of 60kb or above that are flanked on both sides by D domains, we find that 61 (35%) are associated with D-D interaction regions that contain more than 1.1-fold higher interaction density in the D-D region compared to neighbouring bins. A further feature of the interaction map, exemplified in Fig. 6, is that E domain TADs are frequently flanked by a zone of relatively reduced interaction, corresponding to the zone of D-E interactions. The relative lack of interaction in this zone compared to neighbouring E-E and D-D interaction zones suggests that, whilst nearby homologous regions interact, there is less heterologous (D-E) interaction suggesting spatial segregation of D and E regions in the nucleus.

**Fig. 6.**
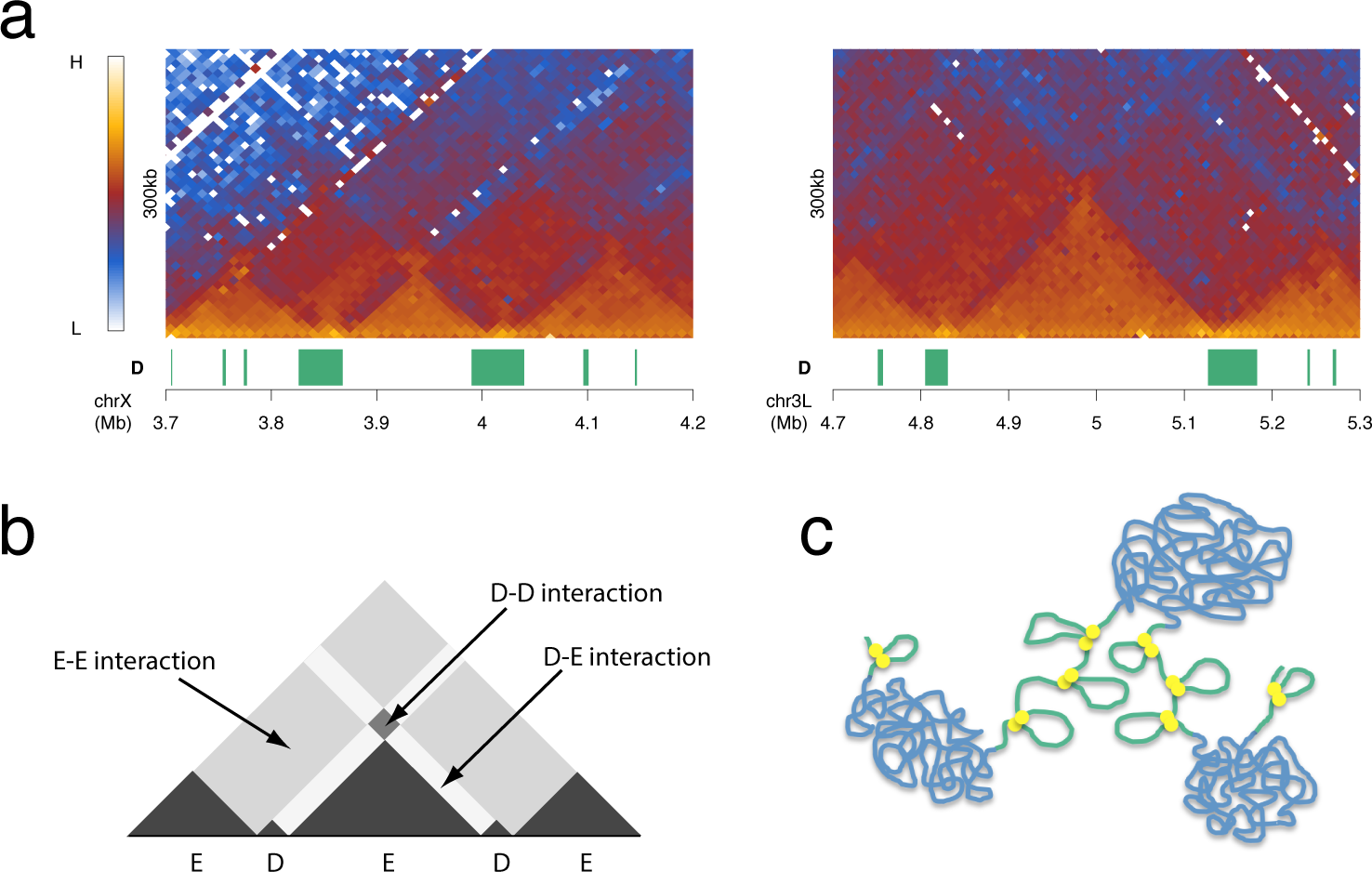
Inter-domain interactions. (a) Kc Hi-C interaction heatmaps at selected D and E domains showing evidence of interaction between D domains. The “D” track shows D domains (green bars). (b) Schematic of interaction heatmap showing inter-domain interactions. (c) Model of chromatin organization into D (green) and E (blue) domains. The two domains types are represented with different chromatin topologies with the E domains condensed and the D domains more open but with defined short-range interactions mediated by insulator complexes (yellow discs). D and E domains are represented as spatially segregated fitting the observation of interaction between domains of the same type.

## Discussion

In this paper we propose a fundamental binary partitioning of the genome based on chromatin state. We identify two domain states based on the level of H3K27me3: Depleted or D domains having little or no H3K27me3 and Enriched or E domains with significant levels of H3K27me3. These two domain states differ markedly in their genomic features. In particular, the D domains are enriched in housekeeping gene TSSs, in regulators of housekeeping genes such as the histone acetyltransferase-containing NSL complex and in a subset of insulator and insulator-related components including BEAF-32, Chromator and Z4/Putzig. In contrast, the E domains are enriched in tissue-specific genes suggesting that the D/E divide may represent a binary subdivision between the constitutive and regulated genome. The organization of D and E domains is strongly associated with the topological architecture of the genome, supporting the idea that the constant TAD organization identified in several studies may be based on prominent TADs being flanked by domains enriched in constitutive gene expression. The prominent TADs correspond to E domains and are flanked by D domains rich in housekeeping genes. This proposed binary subdivision on the basis of D and E domains is likely to be related to the long-observed binary subdivision of the *Drosophila* genome based on the banding pattern of polytene chromosomes with D domains, rich in inter-band-specific proteins such as Chromator, Z4/Putzig and MRG15, representing inter-bands and E domains representing chromosomal bands.

Although we propose a constant partitioning of the genome based on the H3K27me3 profile, the level of H3K27me3 in the E domains is dynamic. Early in development H3K27me3 is focused on Pc target genes, whereas later in development H3K27me3 is at moderate/high levels throughout E domains and this is particularly apparent in the profile from purified germline cells from the adult testis. Increased spreading of the H3K27me3 mark during development has been noted in mammalian genomes [40]. There is also a clear relationship between the widespread H3K27me3 we observe later in development and the profile of H3K27me2, which shows pervasive binding in E domains (except for Pc target regions) [27]. Inhibition of PRC2 histone methylase leads to loss of both H3K27me2 and me3 and is accompanied by a widespread increase in transcriptional activity not just at Pc target genes [27]. Pervasive H3K27me2 is proposed to be governed by opposing roaming activities of PRC2 and the UTX demethylase complexes. This mechanism, together with an active role in widespread gene repression, may also apply to the H3K27me3 in E domains.

The striking correspondence of the D domains, defined by chromatin state, with the regions flanking prominent TADs suggests that the genomic distribution of chromatin state domains may have a key role in establishing the genome interaction architecture. What are the features of D domains that could enable them to function as boundary regions or inter-TADs? Although very low levels of H3K27me3 define D domains, this histone mark seems unlikely to be the key functional property since it is dynamic, in contrast to the TAD organization, which appears to be constant. The chromatin state characteristics of D domains include little or no H3K27me2 and enrichment for active marks such as H3K36me3 and H4K16ac. The major enzyme responsible for acetylation of H4K16 is MOF [35], a component of the NSL complex that regulates housekeeping genes [31, 32] and D domains are enriched in the NSL component MBD-R2. H4K16ac is a candidate for a modification with an effect on genome architecture as it has been demonstrated to affect chromatin structure [41]. It is interesting that HAT complexes and HDACs (particularly HDAC3 which targets H4K16ac [34]) are both enriched in D domains, however both are associated with gene expression in a mechanism whereby the H3K36me3 mark, linked with elongating PolII, recruits HDACs to inhibit inappropriate initiation within active transcription units [42–44]. A further modification that may be relevant to an architectural role of D domains is H3S10 phosphorylation mediated by JIL-1 kinase, which is recruited to polytene chromosome interband regions by a complex containing Chromator and Z4/Putzig [39]. Mutations in JIL-1 kinase, Chromator and Z4/Putzig all result in disruption to polytene chromosome structure [38, 39, 45–47]. Overall, we envisage that housekeeping transcription factors and a subset of insulator components recruit chromatin modification complexes leading to the establishment of the D chromatin state. The observed TAD organization could be dependent on the different properties of the D and E chromatin states. As suggested and modelled by Ulianov et al. [21], TAD organization can be generated based on two domain states with differing chromatin aggregation properties. We have shown that D and E domains differ in their interaction properties (Fig. 5) with E domains showing a rather even high density of interactions across the domain, consistent with condensed chromatin, and D domains being characterised by shorter range interactions. Differential interaction properties are also supported by the observation of decreased interaction between neighbouring D and E regions and increased interaction between nearby D domains (Fig. 6), suggesting spatial segregation of the two domain states.

Insulator proteins have been proposed to be key mediators of both genome architecture and chromatin state domains. Our analysis has implications for insulator complex function since we do not find insulator protein binding to be strongly associated with the D/E chromatin state boundaries. The binding of a subset of insulator components is enriched in D domains but these components bind throughout the D domains and are strongly associated with the TSSs of housekeeping genes, not with domain boundaries. This suggests that insulator components, at least in the BEAF-32/Chromator/CP190 context, may be more directly associated with transcriptional regulation than with chromatin state boundary formation. BEAF-32 has been linked to transcription and in *BEAF-32* mutants many BEAF-32-associated genes show reduced expression [26]. In addition, Chromator acts as a transcriptional activator with specificity for housekeeping promoters [48]. We suggest that at least part of the contribution of BEAF-32, Chromator and CP190 to genome architecture may be indirect as they may act to establish the chromatin state in D domains, for example by recruiting chromatin modifiers such as JIL-1 kinase, enabling D domains to act as boundary regions flanking TADs. As BEAF-32/Chromator/CP190 complexes mediate DNA interactions [37] and D domains are characterised by short-range looping, the insulator complexes may be mediating loop formation within D domains potentially forming promoter/promoter or promoter/enhancer contacts. The idea that insulator complexes within D domains are primarily involved with transcriptional regulation leaves open the question of what actually determines the location of the D/E boundaries. We speculate that the location of the chromatin state boundaries may be specified, not by insulator complexes precisely at the boundary, but by the range of action of chromatin modification complexes that are recruited to D domains by the insulator complexes and other transcription factors.

Despite a considerable body of evidence from studies on the mammalian genome linking the insulator protein CTCF with the domain organization of the genome, we find little association between CTCF binding and the D/E domain organization defined by H3K27me3 levels. Here our studies agree with Ulianov et al. [21], where HiC experiments on several *Drosophila* cell lines found that CTCF was only weakly enriched at regions separating TADs, and with Sexton et al. [9] who found in a genome-wide 3C analysis of embryo chromatin that Chromator is considerably more strongly associated with topological boundaries than CTCF. On the other hand, CTCF is clearly associated with domain architecture as it flanks Pc-regulated domains in Hox complexes [19, 49, 50] and mutating CTCF sites disrupts these domains [19]. Sexton et al. [9] find that CTCF is preferentially associated with borders of Pc domains. This suggests that, at least in *Drosophila*, CTCF may mediate the formation of a specialized set of domains distinct from the more general D/E domain organization of the genome.

## Conclusion

We propose a binary partitioning of the genome based on chromatin state that 454provides a basis for the organization of chromatin into TADs and that reveals an architectural distinction between the organization of constitutive and regulated genes.

## Methods

### ChIP-array on Primary Spermatocytes

*Fly stocks*: For sorting YFP^+^ primary spermatocytes, we used homozygous males of the YFP-tagged protein-trap line *heph^CPTI002406^* [51] which is homozygous fertile.

*Testis dissections, cell extraction and fixation*: Testes were dissected in ice-cold Schneider's medium (supplemented with 10% fetal calf serum) and incubated with collagenase (5 mg ml^−1^, Sigma-Aldrich C8051) plus protease inhibitors (Sigma-Aldrich P8340) in medium for 5 min at room temperature. After washing in medium, cells were extracted by gently pipetting for 5 min in 100 µl medium, using a P200 tip (Rainin RT-200F; with 1.5 mm of the tip cut off to increase the diameter of the opening) and fixed by adding an equal volume of 2% formaldehyde (Sigma-Aldrich F8775) in PBS. Cells were mixed thoroughly and incubated for 10 min at 23°C in an Eppendorf Thermomixer at 700 rpm. Fixation was stopped by adding 400 µl ice-cold medium and placing the sample on ice. The sample was centrifuged in a swing-out rotor at 1,000 *g* for 5 min at 4°C and the pellet snap frozen in liquid N 2 prior to storage at −80°C. A total of 1,000 testes were dissected for each ChIP-array replicate. Testes were dissected in batches of 100 and, for each batch, the time from the start of dissection until fixation was approximately 1 hr.

*FACS*: Aliquots of extracted cells stored at −80°C were thawed, combined in PBS/0.01% Triton X-100 and passed through a 50 µm filter (Partec 04-004-2327). Cells were sorted using a 100 µm nozzle on a MoFlo FACS machine (Beckman Coulter) equipped with a 488 nm argon laser (100 mW). Cells were sorted into a microfuge tube containing 700 µl PBS/0.01% Triton X-100. Events were triggered on forward scatter and YFP^+^ events were sorted using the gating strategy described in Fig. S2. Data was acquired and analysed using Summit software (Beckman Coulter).

*ChIP on sorted cells*: Sorted cells were centrifuged in a swing-out rotor at 4,000 *g* for 15 min at 4°C, transferred to a thin-walled 0.5 ml microfuge tube (Axygen PCR-05-C), re-centrifuged and then resuspended in 130 µl Lysis Buffer (17 mM Tris.HCl (pH 8), 3.4 mM EDTA.Na_2_, 0.34% SDS) containing protease inhibitors (Sigma-Aldrich P8340). The lysate was sonicated for 5 cycles at high setting using a Diagenode Bioruptor (1 cycle is 30 s ON and 30 s OFF). After sonication, the sample was centrifuged at 16,000 *g* for 15 min at 4°C, the chromatin-containing supernatant transferred to a fresh microfuge tube and 70 µl RIPA buffer (36.7 mM Tris.HCl (pH 8), 2.5 mM EDTA.Na 2, 0.01% SDS, 2.46% Triton X-100, 374 mM NaCl) containing protease inhibitors added to the chromatin sample. The ChIP reaction, washes and DNA purification were performed as in Dahl and Collas [52, 53]. In brief, magnetic beads were coated with 2.4 µg of rabbit anti-H3K27me3 antibody (Millipore 07-449) and incubated overnight in a volume of 100 µl with chromatin from ∼100,000 YFP^+^ sorted cells. Beads were washed, chromatin eluted, RNA and proteins digested, the DNA purified by phenol/chloroform extraction followed by ethanol precipitation using linear acrylamide as carrier and resuspended in 10 µl PCR grade water. Approximately 5 µl of chromatin was retained as input and purified alongside the ChIP sample.

*Amplification and labelling of ChIP DNA*: ChIP and input DNA were amplified using the GenomePlex Single Cell Whole Genome Amplification Kit (Sigma-Aldrich WGA4) following the manufacturer's instructions from the library preparation stage. Approximately 150 pg of DNA was used for amplification. Samples were amplified for 22 cycles and amplified DNA was purified using a QIAquick PCR Purification Kit (Qiagen). The amplified ChIP and input DNA (2 µg each) were labelled with Cy5 and Cy3 using the BioPrime DNA Labeling Kit (Invitrogen) in the presence of Cy3- or Cy5-dCTP (GE Healthcare) and hybridised onto Nimblegen ChIP-chip 2.1M Whole-Genome Tiling Arrays according to the manufacturer's instructions.

*Microarray data processing*: We performed two biological replicates with a Cy3/Cy5 dye swap. Input chromatin was used as the reference to determine ChIP enrichment. Arrays were scanned and the images processed using NimbleScan software to generate raw data (*.pair) files for each channel. Loess spatial correction was performed using the NimbleScan software and an in-house R script was used to generate normalised log_2_ ChIP/input ratio (*.sgr) files. For each array the median intensity per channel was scaled to 500 then quantile normalisation was performed across all channels. The normalised ratio scores for both arrays were averaged then smoothed by computing the mean score per 1 kb tiling window. The resulting *.bedgraph file was visually examined using the Integrated Genome Browser [54].

### HiC

HiC protocol was based on the methods described in [6, 36, 55]. *Cell Collection*: Kc167 cells (obtained from the *Drosophila* Genomics Resource Center) were cultured in 10cm Petri plates in Schneider’s medium supplemented with 5% fetal calf serum and antibiotics at 25°C. Cells from 6 plates were harvested into sterile 50 ml centrifuge tubes. The cells were collected by centrifugation at 1,200 rpm for 5 min at 4°C then re-suspended in 10 ml fresh medium and the cell concentration determined using a haemocytometer. 1 x10^8^ cells were resuspended to a total volume of 45 ml in fresh medium, fixed by the addition of 1.25 ml 37% formaldehyde solution and incubated with gentle shaking on a platform shaker at room temperature for 10 min. The reaction was stopped by adding 2.5 ml 2.5M glycine and further incubation at room temperature for 5 min followed by 15 min on ice. The cross-linked cells were divided into 4 x 15 ml falcon tubes (∼12.3 ml per tube) and collected by centrifugation at 1,500 rpm for 10 min at 4°C. The supernatant was discarded and the fixed cells were flash frozen in liquid N_2_ then stored at -80°C.

*Restriction enzyme digestion*: Cells were thawed on ice (∼2.0 x10^7^ cells) and 500 μl of lysis buffer (10 mM Tris-Cl, pH 8.0; 10 mM NaCl, 0.2% Igepal CA360) and 50 μl of protease inhibitor cocktail (Sigma) were added. Cells were lysed with 15 strokes of a Dounce homogeniser using the tight pestle (Pestle A). The cell suspension was transferred to a microcentrifuge tube and centrifuged at 5,000 rpm at room temperature for 5 min. The supernatant was discarded, the pellet washed twice with 0.5 ml of ice-cold 1.20x NEBuffer 3 (5,000 rpm, 5 min at room temperature) and the pellet resuspended in 500 μl 551 of ice-cold 1.20x NEBuffer 3. 7.5 μl of 20% SDS was added and the sample incubated at 37°C with shaking at 900 rpm for 1h, 50 μl of 20% Triton X-100 was added and incubation continued at 37°C, 900 rpm for 1h. A 50 μl aliquot was taken and stored at -20 °C to serve as undigested control. 45 μl of 1.20x NEBuffer 3 was then added to the remaining lysate followed by 400 U (8 μl) of 50U/μl DpnII and the digestion reaction was incubated at 37°C with shaking at 900 rpm overnight.

*Blunt ending fragments with Klenow enzyme*: The restriction enzyme reaction was incubated at 65°C for 20 min to inactivate DpnII, then dispensed into five 100μl aliquots in fresh microfuge tubes. Tube 1 was designated as a 3C control and Tubes 2-5 were used as Hi-C samples. To the Hi-C samples 0.35 μl each of 10 mM dCTP, dGTP and dTTP, 8.75 µl of 0.4 mM biotin-14-dATP and 5 µl of 5U/µl Klenow enzyme (NEB M210S) were added followed by incubation at 37°C for 45 min.

*Ligation*: 200 μl of 10 mM Tris-HCl/1% SDS was added to each blunt-ended reaction and the 3C control tube, followed by transfer to a fresh 50 ml tube containing 700 µl 10x T4 DNA Ligase Buffer (NEB) and 5894 µl molecular biology grade water. Then 106 μl 20% Triton X-100 was added and the mixture incubated for 1 h at 37°C with gentle shaking. 10,000U of DNA ligase (NEB, M0202L) was added and then incubated for 18h at 16° C followed by 30 min at room temperature.

*De-crosslinking and DNA purification*: 30 μl of 10 mg/ml Proteinase K (300 μg final) was added to the reaction and incubated at 65° C overnight. 10 μl of 30 mg/ml RNase A (Sigma) was added and the reaction was incubated at 37° C for 45 min. 7 ml phenol-chloroform-isoamyl alcohol (PCI) was added and mixed vigorously by vortexing. The mixture was transferred to a 50 ml Phase Lock Gel tube (5 Prime, 2302860) and centrifuged at 2,200 g for 15 min at room temperature. The aqueous phase was transferred to a fresh 50 ml tube and 7 ml of distilled water, 1.5 ml 3M NaAcetate pH 5.2, and 35 ml 100% ice-cold ethanol were added. The mixture was incubated at -80° C for 1 h then centrifuged at 2,200 g, 4° C, for 45 minutes. The pellet was washed with 10 ml 70% ethanol, centrifuged at 2,200 g at 4°C for 15 min and the pellet dried at room temperature for 10-15 min. The pellet was then re-suspended in 150 μl of 10 mM Tris-HCl pH 8.0 and allowed to dissolve overnight at 4°C. The purified DNA was then transferred to a microcentrifuge tube and stored at -20°C.

*Re-purification of DNA*: 150 μl of 10 mM Tris-HCl pH 8.0 was added to the dissolved pellet followed by 300 μl Phenol-ChIoroform. These were then mixed and centrifuged at 13,200 rpm for 5 min at room temperature in a Phase Lock tube. Phenol-ChIoroform extraction was then repeated on the aqueous phase and DNA was precipitated by adding 30 µl 3M NaAcetate, pH 5.2 and 2.5 volumes of ice-cold 100% ethanol. The mixture was incubated at -80° C for at least 45 min then centrifuged for 20 min at 13,200 rpm at 4° C. The pellet was washed with 0.5 ml of ice-cold 70% ethanol and air-dried briefly. The pellet was then resuspended in 25 μl TLE Buffer (10 mM Tris-Cl pH 8.0, 0.1 mM EDTA). The Hi-C DNA preparations were pooled into a single tube 24 and the concentration measured using a Qubit photometer.

*Removal of biotin from unligated ends and shearing of Hi-C DNA*: For every 5 μg of Hi-C DNA preparation, the following were added: 1 µl 10 mg/ml BSA, 10 µl 10X NEBuffer 2, 1 µl 10mM dGTP, 1.67 µl (5U) T4 DNA polymerase and water to a total volume of 100 µl. The mixture was incubated at 12°C for 2 hr and the reaction stopped by adding 2 µl 0.5 M EDTA pH 8.0. DNA was then purified by adding 1 volume of Phenol-ChIoroform and centrifuging at 13,200 rpm for 5 min at room temperature in a Phase Lock tube. The DNA in the aqueous phase was precipitated by adding 10 μl 3M NaAcetate, pH 5.2 and 2.5 volumes ice-cold 100% ethanol. The mixture was incubated at -80° C for at least 30 min then centrifuged for 20 min at 13,200 rpm at 4°C. The pellet was washed with 0.5 ml of ice-cold 70% ethanol, then dissolved and pooled in a total volume of 100 μl of water. The Hi-C DNA preparation was sheared at 4°C for 20 min in 30 sec pulses at low setting using a Bioruptor sonicator (Diagenode).

*Library Preparation*: 1.5 μg sheared DNA was end repaired, poly-A tailed and adapters added using the TruSeq Kit (Illumina). The adapter ligated Hi-C library was run on a 1.5% agarose gel in 0.5X TBE at 50V for 2 h. The gel was stained with Sybr Gold and DNA fragments ∼250 to 750 bp in size were excised and purified using a Qiagen Gel Extraction Kit. The DNA was eluted with 50 µl of TLE buffer and the final volume adjusted to 300 µl with water. Biotin-tagged size-selected Hi-C DNA was purified using Dynabeads MyOne Streptavidin Beads (Invitrogen). 150 µl of resuspended beads were prepared according to the manufacturer’s instructions. These were then re-suspended in 300 µl 2x Binding Buffer (10 mM Tris-HCl pH 8.0, 1 mM EDTA, 2 M NaCl) and combined with 300 μl of repaired Hi-C DNA. The biotin-labelled Hi-C DNA was incubated with the streptavidin beads at room temperature for 15 min on a rotator. The beads were then washed in 400 µl 1x Binding Buffer followed by a single wash with 100µl 1x ligase buffer. The beads were then re-suspended in 50µl 1x ligase buffer and submitted to the Eastern Sequence and Informatics Hub (EASIH, Cambridge) for library preparation and 76-bp paired-end sequencing on an Illumina GAIIx sequencer.

### HiC data and segments

Paired-end HiC reads were aligned against *Drosophila* genome BDGB release 5 with HiCUP (v0.5.7) [56]. Read counts/filtering are given in Table S1. Interactions were binned at 10 kb resolution. Contact matrices were normalised using the GOTHiC_1.6.0 R package [57]. HiC segments were identified from the normalised contact matrices using the HiCseg R package [58], with 10% of the lowest linear portion of log-likelihood segment borders removed.

### Gene Classes and H3K27me3 developmental profile

Housekeeping genes (4,091 genes) were derived from FlyAtlas [59] selecting genes expressed in all tissues (4 present calls), spermatogenesis genes (1,428 genes) were from Chen et al. [60] (GEO accession GSE28728) selecting genes down-regulated 4-fold or more in *aly* mutant testes, not-expressed-in-testis (2,432 genes) were from FlyAtlas selecting genes with zero calls in testis and >4 positive calls across other tissues and Polycomb_target genes (359 genes) were from Kwong et al. [61]. Genome annotation was Flybase Release 5.57 and the “all” class includes all genes in the euchromatic genome (13,832 TSSs). Raw ChIP-chip files of H3K27me3 in *Drosophila* at different time points of development were downloaded from GEO GSE15423. The data was quantile normalised for each time point and average scores across replicates calculated. Median scores for all gene TSSs (+/- 500bp) were calculated for each developmental stage. For the relative distance analysis, the mean scores of housekeeping, Pc and spermatogenesis TSSs were calculated for each stage. The distance between housekeeping and Pc class means was set to 1 and the relative distance of the spermatogenesis class mean to the housekeeper mean was calculated.

### H3K27me3 Hidden Markov Model (HMM)

Normalised Spermatocyte H3K27me3 euchromatin oligo scores were binned at 1 kb resolution taking the mean score per bin. A HMM using normal distribution was fitted for two states using the RHmm R package [62] for each chomosome. This divided the data into D (depleted in H3K27me3) and E (enriched in H3K27me3) bins. Adjacent bins with the same HMM state were then combined into regions. Gaps in the genome for which there were no oligo probes were closed if the same state was present on both sides.

### Binding data

ChIP-chip.cel files were downloaded from ModENCODE [63] for Chromator (277), CTCF (908), GAF (2568) and Jil1 (3037) and processed with Ringo [64] using a half window of 300 bp, minProbes 8 and max gap 200 bp. Binding regions were called using a False Discovery Rate (FDR) of 1%. ChIP-seq fastq data was downloaded from GEO for BEAF32 (GSM1535963), CP190 (GSM1535980), Z4/Putzig (GSM1536022) [65] and Su(Hw) (GSM762839) [66]. Reads were mapped to *Drosophila* genome BDGB release 5 using Bowtie with -m 1 option. ChIP-seq peaks were called with macs2 2.1.0 (https://github.com/taoliu/MACS) [67] using unique tags and a q-value threshold of 1e-6 and otherwise the default parameters. Data sources are given in Table S2.

### Fraction of genes in class

Gene expression score data for 25 *Drosophila* cell lines were downloaded from Supplemental Table S-3 of Cherbas et al. [68]. These gene scores derived from whole-genome tiling microarrays and represent the normalised maximum score for all exons included in that gene. Genes were assigned to D and E regions based on the location of the TSS. As suggested by Cherbas et al. we selected a threshold score of 300 to distinguish the expressed from unexpressed genes. For each gene we counted in how many cell lines it had an expression score > 300, and calculated the fractions of expressed genes in D and E.

### Interaction Fractions

HiC interactions were binned at 5kb resolution and normalised using GOTHiC. Interactions for each 5 kb genomic bin up to 1 Mb distance were collected, excluding genomic coordinates located within 1 Mb of the chromosome ends. The sum of the interactions for each bin were set to 1 and fractions for close (5-50kb, thus ignoring interactions which are within 5kb of each other) and far (50-500kb) calculated. Bins with no interactions and bins with extreme high interaction sum (> 97.5^th^ percentile) were excluded. The ratio is given as fraction close/fraction far.

### Declarations

#### Ethics approval and consent to participate

n/a

#### Consent for publication

n/a

#### Availability of data and material

The datasets generated during and/or analysed during the current study are available at GEO (Accession number GSE85504) 708

#### Authors’ Contributions

SE, JPM, SWC, SR and RW conceived and supervised experiments. SE and JPM performed the experiments. SE, BF, BU, RF and RW analysed data. SE, BF, JPM, SR and RW wrote the paper. All authors read and approved the final manuscript.

## Acknowledgements

n/a

## Additional Files

Additional File 1: Fig. S1. The comparison of Kc cell and embryo interaction maps. (PDF 1.2MB)

Additional File 2: Fig. S2. The purification of primary spermatocytes by FACS. (PDF 3.5MB)

Additional File 3: Table S1 HiC/HiCUP read counts and filtering. (XLSX 11KB)

Additional File 4: Table S2 Data sources. (XLSX 11KB)

